# Quantification of protein mobility and associated reshuffling of cytoplasm during chemical fixation

**DOI:** 10.1101/410670

**Authors:** Jan Huebinger, Jessica Spindler, Kristin J. Holl, Björn Koos

## Abstract

To understand cellular functionalities, it is essential to unravel spatio-temporal patterns of molecular distributions and interactions within living cells. The technological progress in fluorescence microscopy now allows in principle to measure these patterns with sufficient spatial resolution. However, high resolution imaging comes along with long acquisition times and high phototoxicity. Physiological live cell imaging is therefore often unfeasible and chemical fixation is employed. However, fixation methods have not been rigorously reviewed to preserve patterns at the resolution at which they can be nowadays imaged. A key parameter for this is the time span until fixation is completed. During this time, cells are under unphysiological conditions and patterns decay. We demonstrate here that formaldehyde fixation takes more than one hour for cytosolic proteins in cultured cells. Associated with this, we found a distinct displacement of proteins and lipids, including their loss from the cells. Other small aldehydes like glyoxal or acrolein showed inferior results. Fixations using glutaraldehyde were faster than four minutes and retained most cytoplasmic proteins. Surprisingly, autofluorescence produced by glutaraldehyde was almost completely antagonized by supplementary addition of formaldehyde without compromising fixation speed. These findings indicate, which cellular processes can actually be reliably imaged after a certain chemical fixation.

## Introduction

Fluorescence microscopy has advanced to allow for the determination of the precise localization of individual molecules in cultured cells down to nanometer precision ^1,2^. Furthermore, it is now possible to image molecular reactions quantitatively and spatially resolved by microspectroscopy or antibody based methods ^3,4^. In principle, this allows to extract invaluable information about cellular functionalities, which are encoded in spatial organization. However, sample preparation methods have not yet co-developed to fully exploit the potential of these methods. Undoubtedly, sample preparation has to preserve the cellular state with at least the precision of the microscopic readout, to avoid artefacts.

Fluorescence microscopy can in principle be performed on living cells. This is optimal to observe cellular dynamics in all cases, where image acquisition is much faster than the process under investigation. However, more sophisticated superresolution and microspectroscopy methods usually require too long acquisition times to image the rapid processes in living cells ^5^ and they are too phototoxic ^6^. Therefore, cells have to be fixed before imaging. It is possible to cryo-fix cells in a close to physiological state for high resolution imaging ^5,7–9^. However, this necessitates specialized equipment and knowledge and is therefore far from being standard yet. More importantly, cryo-arrest cannot be used for antibody-based approaches, such as immunohistochemistry or *in situ* proximity ligation assay (PLA)^4^, because it does not allow for permeabilisation of the cells. For these reasons, cells are usually chemically fixed before high-resolution or functional imaging. The methods for chemical fixation have been developed decades ago and their impact on the structure of cells has been studied to quite some extent by transmission electron microscopy ^8^. However, cellular transmission microscopy provides mainly structural information about lipid-bilayer enclosed organelles and macromolecular complexes, while single molecules are usually not detectable. Fluorescence microscopy yields complementary information. Distribution of molecules or even their interactions can be mapped over the cells, whereas the surrounding structure of the cell remains invisible. While immunofluorescence has been used for decades to assign molecular localization to certain cellular organelles, the last 20-30 years have seen an enormous improvement of fluorescence microscopy techniques. However, the possibilities to image single fluorescent molecules, quantify distributions of molecules and map their interactions within cells ^1–4^, also raises the requirements for fixation methods substantially. Obviously, any changes introduced during fixation will ultimately lead to an incorrect image of the living cell. It is therefore crucial to know, if and how molecules are rearranged upon chemical fixation. By comparing live cell imaging with cells after fixation some large-scale rearrangements may be detected and certain fixation protocols may thus be identified as inappropriate (e.g. ^10,11^). However, fixation is necessary exactly in those cases, where artefact-free live-cell imaging is not possible. This prohibits this kind of comparison for high resolution imaging. Yet, the duration of chemical fixation can be informative here. This duration is utterly important since cells are in a non-physiological, partially-fixed state until fixation is completed. A change in reaction parameters, can alter cellular reactions during this phase. For a reaction in thermodynamic equilibrium, one can try to keep the reaction parameters (such as pH, temperature, pressure) constant, to keep it balanced. Unfortunately, cells usually maintain their spatial organization away from equilibrium by permanent energy consumption. This results in spatial patterns which depend on various reaction rates and the movement of molecules involved (see e.g. ^12,13^). It cannot be expected that all of the reaction rates and movements are slowed down during fixation in a balanced way and physiological patterns are therefore most likely not maintained. Thus, fixation needs to be fast compared to the decay of the patterns, which is ultimately limited by the molecular movements involved. However, the only systematic investigation on molecular species measures fixation of membrane molecules and concludes that not all of them can be fixed by chemical fixation and fixation time is at least 30 min ^14^. These results preclude to analyse lateral distributions of membrane molecules after fixation. However, membrane molecules are a special case, because of their hydrophobic environment, which might protect them against hydrophilic fixation agents. It is therefore not clear to which extent these results are informative for non-membrane molecules.

Here, we measured fixation times directly on cytosolic proteins by consecutive bleaching of the fluorescent protein mCitrine in the cytoplasm of HeLa cells during the process of aldehyde-fixation. Chemical fixation can in principle also be done by immersing cells in organic solvents, e.g. acetone, ethanol or methanol. Organic solvents denature and coagulate or extract cellular molecules and hence lead to more severe rearrangements in the cytoplasm ^10,15^. Thus, aldehyde fixation is usually the method of choice for fluorescence microscopy. We found that fixation times greatly vary from less than four to more than 60 minutes, depending on the aldehydes used. We further observed blebbing of plasma membrane lipids following all tested aldehyde fixations. Using most tested aldehydes, this was associated with loss of unfixed cytosolic protein from these blebs upon their disintegration. The loss of protein was not measurable upon fixation with glutaraldehyde, which was also the fastest fixation. Unexpectedly, addition of formaldehyde repressed glutaraldehyde-induced autofluorescence almost completely, when added to 1-2% glutaraldehyde, without significantly compromising fixation speed or retention of cytosolic proteins. It seems therefore optimal to fix cells with at least 1% glutaraldehyde in combination with formaldehyde. The speed of the molecules involved should however always be considered, given the fact that fixation times in the order of minutes are not short compared to most cellular processes.

These results should thus give a good basis for researchers to judge, if and which chemical fixation can be applied to image a certain cellular process.

## Results

To measure the time after chemical fixation until movement of cytoplasmic proteins is halted, we performed fluorescence recovery after photobleaching (FRAP) experiments. For this, HeLa cells were transfected with the cytoplasmic fluorescent protein mCitrine and a circular area with 2-μm diameter within the cells was repeatedly bleached at defined time points after the onset of fixation (Fig. S1). Fluorescence intensity increase in the spot after bleaching is a measure for diffusion of unfixed cytosolic mCitrine back into the bleached area. However, dark state recovery of the fluorophore can also contribute to the observed fluorescence recovery^16^. To control for the latter, we performed the same bleaching protocol with cells that were fixed for 1 h (Fig. S2). If the respective recoveries do not differ from this control, no more fixation took place within the following hour. The first time point for which this is the case can therefore be considered as the time when fixation was completed. Using this method, we found that fixation with 4% formaldehyde (FA), which is the standard fixation for fluorescence light microscopy, takes more than 20 min (Fig. 1A; Fig. S2). Radial analysis of full recovery curves obtained after more than 1 h showed some residual diffusion of mCitrine, indicating that even the controls showed not only dark state recovery in this case (Fig. S3). During the first 20 min of exposure to 3% of the small aldehyde glyoxal, diffusion closed the bleached spot before the first post-bleaching image could be acquired, indicating that there is no fixation of cytoplasmic protein within this time. After more than 60 min the onset of fixation was measurable (Fig. S4). This slow fixation is potentially due to the poor membrane permeability of glyoxal ^17^. In contrast, glutaraldehyde (GA) fixation, which is often used for electron microscopy, fixed cytoplasmic proteins within 4 min (Fig. 1A; Fig. S2). In this case, no residual movement was measurable in the controls that were fixed for 1 h, demonstrating that the controls show only fluorescence dark state recovery in this case (Fig. S3). Glutaraldehyde is not routinely used for fluorescence light microscopy, since it is notorious to cause high autofluorescence. Quantification of autofluorescence showed a nearly 400-fold increase in red fluorescence (excitation 535–550 nm/ emission 570–625 nm), a 40-fold increase in green fluorescence (ex. 360-370 nm/ em. 420-470 nm), but no increase in blue fluorescence (ex. 360-370 nm/ em. 420-470 nm) (Fig. 1B). This autofluorescence can be quenched completely in the green and to approximately 5-fold in the red fluorescence channel by application of first 100 mM NH_4_Cl for 40 min and subsequently 5 mg/mL NaBH_4_ for 2 h (Fig. S5). However, this procedure is relatively time-consuming, and its influence on the structure of cells is largely unknown. Interestingly, fixation by a combination of GA and FA (but not by a consecutive addition of the two (Fig. S5)) results in much lower autofluorescence than GA alone, even if the same concentration of GA is used (Fig. 1B). Combinations of 4% FA with GA at 1-2% gave fixation times of < 4 min, similar to 2% GA alone (Fig. 1A).

**Figure 1:**
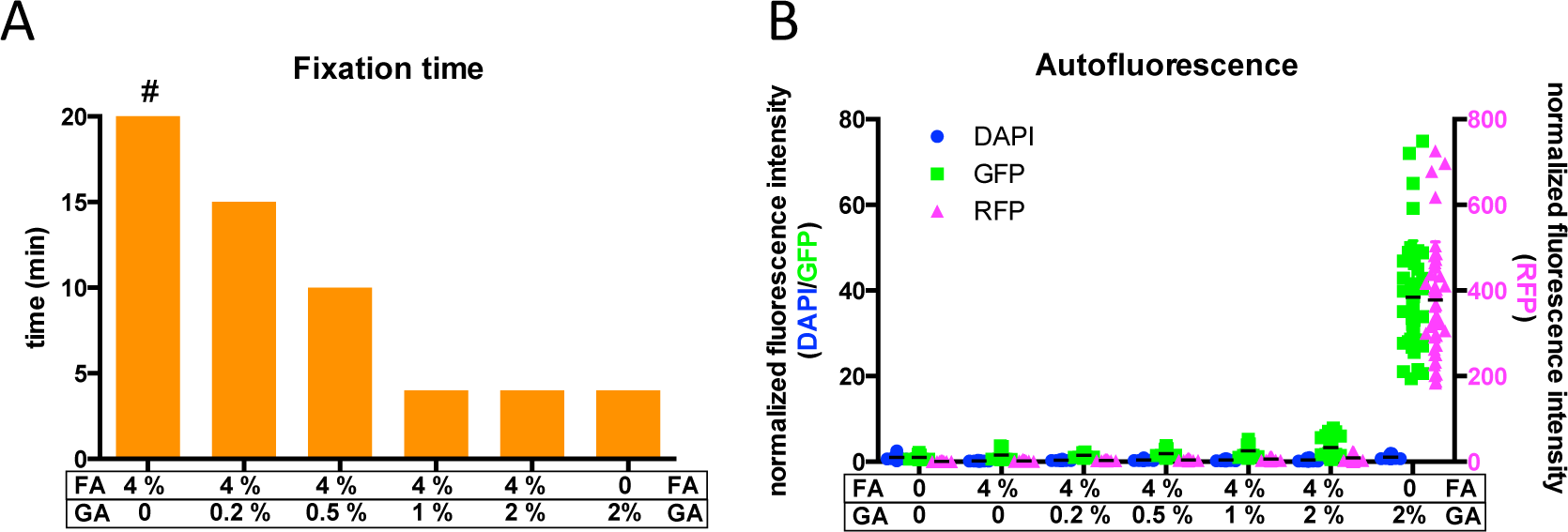
Fixation time of cytoplasmic protein and development of autofluorescence by aldehyde-fixation in HeLa cells. **A)** Fixation time for cultured HeLa cells as determined by consecutive bleaching and fluorescence recovery of mCitrine during chemical fixation using the indicated aldehyde concentrations (Fig. S1-S2). Depicted are the time points after which no further diffusion of mCitrine was observed except **#**, where diffusion was measurable during the whole course of the 20-min experiment after fixation and also in separate experiments after more than 60 min (Figure S2-S3). **B)** Increase in autofluorescence of HeLa cells upon fixation with the indicated concentrations of aldehydes in 3 different fluorescence channels corresponding to blue (DAPI; excitation 360-370 nm/ emission 420-470 nm), green (GFP; ex. 360-370 nm/ em. 420-470 nm) and red (RFP; ex. 535–550 nm/ em. 570–625 nm) fluorescence measured by widefield microscopy normalized to living cells. Data is shown for single cells (colored symbols) and mean (black lines). All autofluorescence measurements in GFP and RFP channels are significantly higher (p<0.001 using student´s t-test) than in living cells. All measurements in DAPI channel, except after fixation with 2% GA alone (n.s.), are significantly lower (p<0.001 using student´s t-test) than in living cells. n=5 independent experiments

Lipids are not fixed at all using aldehyde fixation ^14^. In agreement with this, fluorescent microscopy of cells stained with the lipid dye DiIC12 showed extensive plasma membrane blebbing during the fixation process as visualized by fluorescence microscopy (Fig. 2; Fig. S6-S8). These blebs disintegrated either spontaneously or latest after treatment with the detergent triton x-100, which is regularly used to permeabilise cells for antibody staining. Blebs were observed during GA, FA, glyoxal and acrolein fixation (Fig. 2A; Fig. 6-8). During FA, glyoxal or acrolein fixation, these blebs were filled with cytoplasmic protein, which eventually got lost upon disintegration of the blebs (Fig. 2B; Fig. S4; Fig. S6-S7). Consequentially, fixation with FA, glyoxal or acrolein induced a loss of unfixed cytoplasmic protein. Upon GA fixation, blebs were free of cytoplasmic protein, indicating that no loss of cytoplasmic protein arose (Fig. 2B; Fig. S8). In addition, a membrane-bound fluorescent protein (mCherry attached to the plasma membrane by the membrane-anchor of kRas) was detected in blebs during FA-fixation, but not during GA-fixation (Fig. S9).

**Figure 2:**
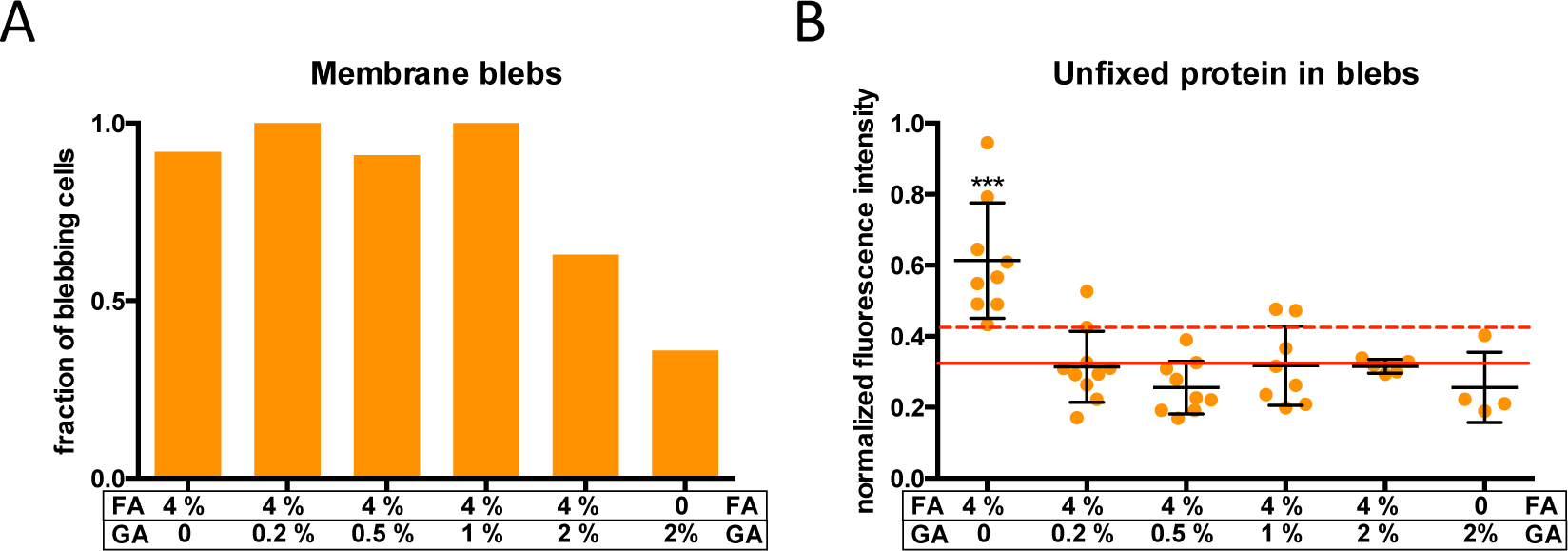
Membrane blebbing and associated loss of cytosolic protein in HeLa cells upon fixation with glutaraldehyde (GA) and formaldehyde (FA). **A)** Depicted is the fraction of HeLa cells that showed membrane blebs after fixation as identified by DiIC12-staining (compare Fig. S6-S8). **B)** Depicted is the mean fluorescence intensity of cytoplasmic EGFP in the plasma membrane blebs normalized to the mean fluorescence intensity of the corresponding cell body for each blebbing cell (orange circles) and the mean +/-s.d. (black lines). The solid red line and the dashed red line indicates mean and the mean + s.d. of the background fluorescence intensity around the cells. ***: p<0.001 vs background fluorescence intensity using student’s t-test; n=6 experiments

Temperature and pH are important reactions parameters. If they are altered during the fixation process, it can be expected that even those physiological reactions that are in thermodynamic equilibrium are changed before fixation is completed. Therefore, all previous measurements in this study were performed by fixing the cells at physiological temperature and pH. However, FA-fixation at 4°C or on ice is quite frequently performed. We therefore performed the same measurements as above on HeLa cells during FA-fixation at 4°C. Similar to FA-fixation at 37°C, cytoplasmic mCitrine was not completely fixed during the first 20 min of fixation. However, we observed blebs only on 20% of the cells. The lower occurrence of blebbing might be due to a phase change in the membrane. This indicates that by lowering the temperature the loss of cytoplasmic protein might be reduced. Yet, since the fixation-time was not faster, the internal state of the cytoplasm is not better preserved. We further tested a recently published protocol utilizing glyoxal in combination with ethanol at pH 4 ^17^. In our hands, fixing cells with this protocol at 37°C led to blebbing in all observed cells accompanied by a complete loss of fluorescent, cytosolic protein (mCherry) and a redistribution of plasma membrane bound protein (mCherry-tkRas) and lipid (DiIC12) to intracellular compartments (Fig. S10).

The previous results suggest fixation with 1% GA in 4% FA at 37°C and physiological pH to achieve the best fixation speed in combination with little autofluorescence. They also show that the fluorescence from expressed proteins (mCitrine, EGFP, mCherry) or lipid dyes (DiIC12) is preserved by this treatment. However, for antibody based methods epitope accessibility is also crucial. We therefore compared fixation in 4% FA with fixation with 1% GA in 4% FA in diffraction-limited immunofluorescence experiments. For most stainings (Rab5, Lamp2, GM130, DAPI, F-actin), which have been previously established and used in FA-fixed samples, the localization to the corresponding organelles (early endosomes, lysosomes, Golgi apparatus, nucleus, actin fibres) was indistinguishable between fixations with 1% GA in 4% FA or FA only. Only staining with an antibody against HSP60 did not show the expected mitochondrial localization, which was observed in FA-fixed samples, in samples fixed in 1% GA in 4% FA exhibited clear but diffuse fluorescence, which could be unspecific binding of the antibody. On the other hand, an antibody against alpha-tubulin gave clearly a complete decoration of microtubules in six out of six experiments after fixation with 1% GA in 4% FA. After fixation with FA alone only dotted stainings along the microtubules were visible, when the same antibody concentrations were used in parallel experiments. Further, testing of an antibody against PTP1b, which we had previously not validated for immunofluorescence, gave only after fixation with 1% GA in 4% FA significant fluorescence signal (Fig. 3). The localization of this ER-bound protein throughout the cell is in agreement with its localization in living cells ^18^. These results show that epitope accessibility is specific for combinations of antibody and fixation method and none of this two fixation methods is generally favourable in this regard.

**Figure 3:**
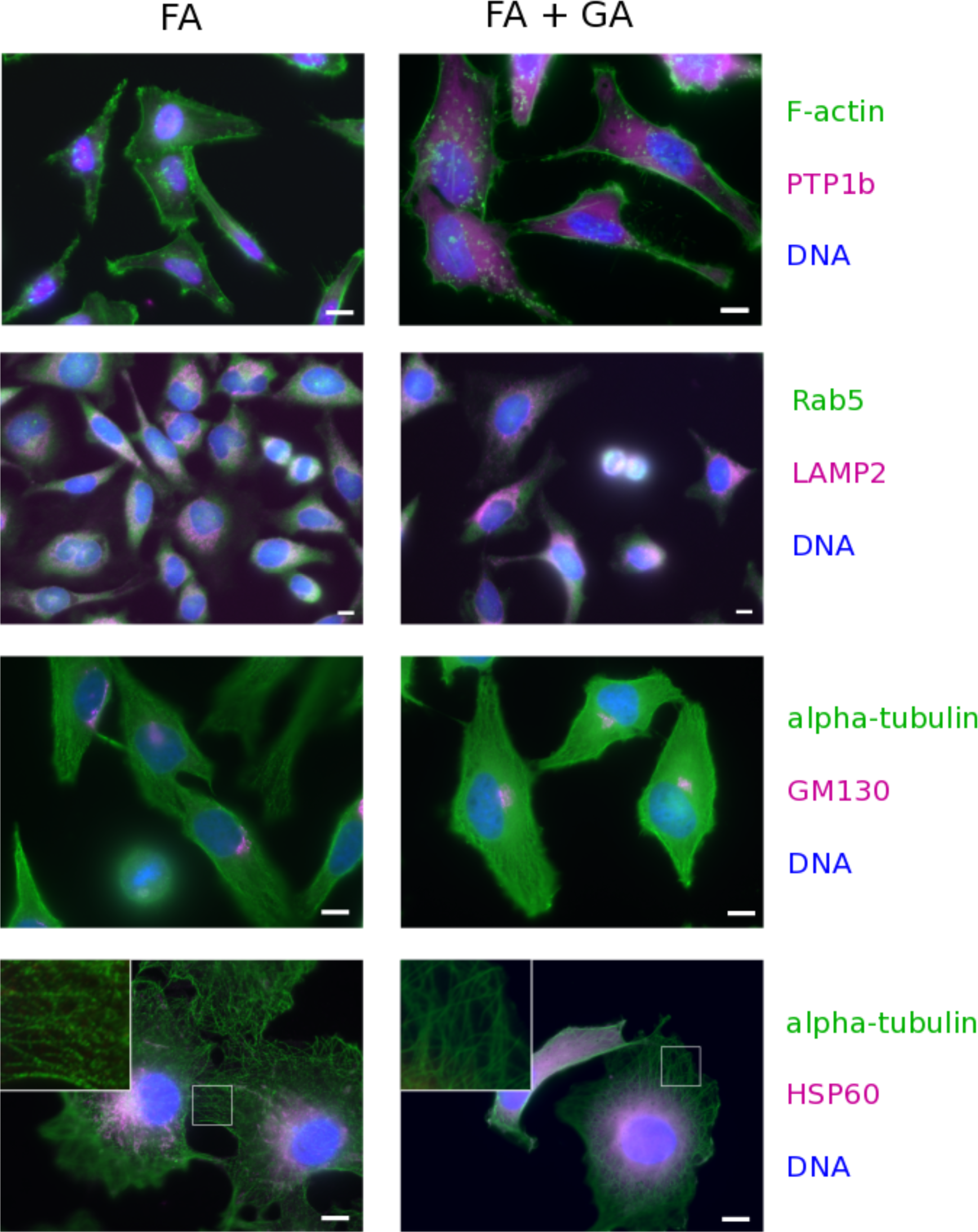
Different fluorescent stainings in cells fixed with formaldehyde (FA) and a combination of glutaraldehyde (GA) and FA. HeLa cells (rows 1-3) or Cos7 cells (4^th^ row) have been fixed with 4% formaldehyde (FA; left column) or 1% GA in 4% FA (FA + GA; right column). The cells have been permeabelised and the molecules indicated on the right of each row in the corresponding colours have been fluorescently labelled. The proteins PTP1b, Rab5, LAMP2, alpha-tubulin and HSP60 have been labelled by indirect immunofluorescence. F-actin has been labelled by Alexa488 coupled to Phalloidin. The DNA has been labelled with the intercalating dye Hoechst 33342. All experiments have been repeated three times and the images shown are representative. The inserts in the last row show magnifications (3.3x) of the boxed area in the corresponding images. Scale bars: 10 μm

## Discussion

The presented results provide an overview on fixation times, autofluorescence, membrane blebbing and loss of cytoplasm during different aldehyde fixations. We have mainly focused on the aldehydes that are most widely used for fixation, FA and GA, and their combinations. Other small aldehydes, acrolein and glyoxal, that are only rarely used for fixation, gave under the tested conditions inferior results to FA and GA. Generally, our results suggest to perform chemical fixation for cytoplasmic proteins by a combination of FA and GA with GA-concentrations of at least 1% at physiological temperature and pH. This fixation method was among the fastest, retaining most of the cytosolic proteins, while producing only very minor autofluorescence, which – for very sensitive measurements – could be even further quenched. Combinations of GA and FA are rarely used for modern fluorescence light microscopy. In these attempts, usually very low (often around 0.1%) concentrations of GA are used to prevent autofluorescence and we show that GA concentrations need to be at least 1% to achieve optimal fixation times. Surprisingly, our results also show that the autofluorescence induced by GA is very much reduced in the presence of FA, which allows for higher concentrations of GA and thereby much faster fixation. It is of note that the fixation-artefacts that we have observed here are very obvious during the fixation process. However, extraction of lipids and proteins as well as rearrangements in their organization would not be self-evident in fixed cells alone, which could lead to false interpretations, if the process of fixation is not completely understood.

We further show that epitope accessibility depends on the combination of antibody and fixation. Most antibodies that are established in FA-fixed samples worked also after fixation with a combination of FA and GA. Yet, this is not always the case and some antibodies only bind after one of the fixations. However, we think that an appropriate fixation, which preserves the cell in a closer to physiological state, should be the primary objective. The antibody should comply with the fixation method not the other way around, whenever possible.

However, based on the measured process and the desired readout, differing fixation methods might be sufficient or even slightly beneficial. Due to the wide variety of fluorescence light microscopy measurements this needs to be assessed on a case-by-case basis. This however necessitates knowledge about the course of fixation. Changing pH or temperature will change the physiological state of the cell and did not result in faster or more efficient fixation. It may therefore only in very special cases be advisable to do so. E.g., if the amount of a cytoplasmic protein is to be quantified in a cell, but the subcellular distribution is not of interest, fixation with 4% FA on ice (but not at 37°C) might be sufficient. This fixation is much slower than fixation with a combination of GA and FA at 37°C, however the autofluorescence in the green to red spectrum should be slightly lower and due to the reduced blebbing at 4°C, loss in cytoplasmic protein is less than at 37°C. Further, if blue fluorophores are used, also fixation with 2% GA can be efficient, since we found aldehyde-induced autofluorescence only in the green and red part of the spectrum. Also, to find out if a protein of interest is located in a certain membrane-bound organelle, even the rather inefficient fixation with formaldehyde at 37°C can be sufficient and this has actually been done in numerous studies over decades. Even though this is not optimal and e.g. a putative cytoplasmic fraction might be underestimated due to extraction of cytoplasmic protein.

Modern microscopy is however able to measure the distribution of cellular macromolecules and even their interaction at much higher resolution. In this context, it is important to note that even the fastest tested fixations happen in the order of minutes, which is slow compared to many cellular processes. Imaging a fixed cell can thereby never be considered as taking a snapshot of a living cell at a distinct point in time, as it is often implied. In addition to a good understanding of the process of fixation, also knowledge about the investigated cellular process itself is necessary to judge, if a certain fixation is actually suited for a particular measurement. Some cellular structures might be in or close to thermodynamic equilibrium. These are relatively uncritical to fix, since they will not change as long as the reaction parameters (e.g. temperature, pressure, pH) are not changed. Yet, they still could be altered, if extraction from the cytoplasm changes concentrations of reaction partners. However, most processes in a living cell are maintained out of equilibrium by permanent energy consumption, as this is an inherent feature of life ^19^. For all of these processes, fixation speed is very critical, since it needs to be faster than the decay of these patterns upon withdrawal of energy. A well investigated example for a spatial regulated pattern in the cytoplasm is signal transduction to the nucleus after sensing of extracellular stimuli at the plasma membrane of cells by receptor molecules. The receptors initiate a cascade of activation events in the cytoplasm, usually by phosphorylation of proteins. These phosphorylated proteins diffuse away from the plasma membrane. However, in the cytoplasm phosphatase activity is dominating, which reverts their phosphorylation ^12^. The phosphorylation of proteins at the plasma membrane (the source) and their diffusion into the dephosphorylating cytoplasm (the sink), creates a gradient of phosphorylation from the cytoplasm into the interior of the cell ^12^. This gradient is dependent on at least three factors: the rates of phosphorylation and de-phosphorylation and the speed of diffusion. Diffusion will obviously be slowed down by fixation, but also phosphorylation and de-phosphorylation are non-equilibrium processes and will be altered during the lethal fixation process ^20^. Since one cannot expect these processes to be slowed in a balanced way, it will alter the appearance of the gradient in the cell after fixation. Cytosolic proteins diffuse with 10-100 μm^2^/s. Therefore, gradients in the cytoplasm can decay in the timeframe of a second or less. This can also be seen from FRAP experiments. If a spot in the cytoplasm of unfixed cells is bleached, this represents an artificial gradient, which is not maintained by energy consumption. This spot disappears within less than a second, which can for example be seen in figure S4A. This is of course an extreme case, since it cannot be expected that upon fixation the energy is immediately depleted, but it might be useful to illustrate the fragility of diffusion-dependent patterns. Thus, it might be not possible to fix such patterns with the available chemical fixations and consequently their direct imaging in fixed cells has – to our knowledge – not been achieved yet. This is of course also true for more complicated diffusion-dependent patterns like Turing patterns, which could also be formed within cells ^12^. On the other hand, structural elements such as some parts of the cytoskeleton have lower turnover rates and are therefore somewhat easier to fix. In conclusion, the results presented here should provide a good basis for researchers to make an informed decision, if and which chemical fixation is suitable to measure a specific cellular process.

## Material & Methods

### Preparation of fixation media

A commercially available phosphate-buffered 4% formaldehyde solution was used to generate the results presented in this study (Histofix, Carl Roth GmbH, Karlsruhe, Germany). However, solutions of formalin diluted to 4% formaldehyde in phosphate buffered solution (PBS) or HEPES-buffered DMEM were also tested. Results were the same for these solutions. Glutaraldehyde (25% stock solution, Sigma-Aldrich Chemie GmbH, Munich, Germany), and acrolein (90% stock solution, Sigma-Aldrich Chemie GmbH, Munich, Germany) were diluted in PBS to the indicated concentrations. Mixtures of GA and FA were obtained by diluting GA (25% stock solution) into the commercially available phosphate-buffered 4% formaldehyde solution (Histofix). Glyoxal was either used as 3% solution in PBS or prepared as suggested in a recent publication ^17^. For the latter case, 2.835 mL ddH2O, 0.739 mL ethanol, 0.313 mL glyoxal solution (40%; Sigma-Aldrich Chemie GmbH, Munich, Germany) and 0.03 mL of acetic acid were mixed and the pH was adjusted to 4 by adding NaOH.

### Preparation of cover slides for cell culture

In order to observe cells during the process of fixation without changing focus or position during addition of the fixative, a self-built flow-through chamber was used ^5^. This necessitated the preparation of special cover slides ^21^. A 10 μm thick double-sided adhesive tape (Modulor GmbH, Berlin, Germany) was cut to the size of a 21 × 26 mm cover slide (Gerhard Menzel GmbH, Braunschweig, Germany). In the centre, a rectangular area with the dimension of the cavity of the aluminium flow chamber was cut out. The adhesive tape was glued on the cover slides (keeping the upper side of the tape covered by the release liner), sterilized in ethanol and washed with sterile H_2_O. Cover slides with adhesive tape facing upwards were placed in sterile 6-well-plates for cell culture (Sarstedt AG & Co, Nümbrecht, Germany).

### Cell Culture and Transfection

HeLa cells (ATCC No. CCL-185) and Cos-7 cells (ATCC No. CRL-1651) were cultured in Dulbecco’s Modified Eagle Medium (DMEM) supplemented with 10% fetal bovine serum (FBS), 100 µg/mL streptomycin, 100 U/mL penicillin, 1% L-Glutamine (200mM), and 1% nonessential amino acids (all PAN-Biotech, Aidenbach, Germany). HeLa Cells were authenticated by Short Tandem Repeat (STR) analysis and did not contain DNA sequences from mouse, rat and hamster (Leibniz Institute DSMZ). Cells were regularly tested for mycoplasma infection using the MycoAlert Mycoplasma detection kit (Lonza, Basel, Switzerland) and cultured at 37°C with 95% air and 5% CO_2_. Cells were plated on the prepared 6-well dishes containing the prepared cover slides or in glass bottom dishes (MatTek Corporation, Ashland, MA, USA) 1-3 days before the experiment. One day before imaging, HeLa cells were transfected using FuGENE™ 6 (Promega, Madison, WI, USA) following the standard protocol of the manufacturer.

### Mounting of the cover slides to a temperature-controlled flow-through chamber

The assembly of the temperature controlled flow-through chamber was performed as previously described in detail ^5,21^. Briefly, the cover slides with cells were removed from the 6-well plates. The release liner of the double-sided adhesive tape was removed and the cover slides were glued to the custom-made flow chamber ^5^. The rectangular part in the middle without adhesive tape was mounted right under the cavity of the flow chamber. Care was taken to keep the cells moist by medium during this process. The temperature control was fixed to the flow-through chamber as previously described. Solution exchanges during cooling and warming were realized with the help of a neMESYS low-pressure syringe-pump (Cetoni GmbH, Korbußen, Germany) that enables a constant flow rate. For live-cell imaging, cells were flushed with HEPES buffered DMEM medium without phenol red and temperature was set to 37 °C. Washing the cells before fixation with PBS, as it is common practice in many fixation protocols, did not yield in significantly different results. For fixation on the microscope, the corresponding fixative was flushed with 3 μL/s and the temperature was set to 37 °C, except for one set of experiments where cells were fixed with FA at 4 °C. Temperature change was timed such that the fixative accessed the cells directly once the temperature reached 4 °C.

### Fluorescence imaging of EGFP, mCherry, mCherry-tkRas and DiIC12

HeLa cells were transfected with plasmids for EGFP only, mCherry only, or EGFP and mCherry-tkRas (mCherry fused to the last 20 amino acids of kRas) one day before the experiment. Labelling of cells with DiIC12 was performed by incubating the cells in 2 μg/mL DiIC12 in FBS-free medium for 2 min and subsequently washing them with HEPES-buffered DMEM medium without phenol red. Cells were imaged on a widefield microscope (Olympus IX-81) with a 63x 1.3 NA oil-immersion objective (Olympus GmbH, Hamburg, Germany). Fluorescence intensity in the blebs was quantified using the software Fiji ^22^.

### Fluorescence Recovery After Photobleaching (FRAP)

FRAP measurements were performed on a Leica SP5 confocal microscope using a 63x 1.4 NA oil-immersion objective (Leica Microsystems CMS GmbH, Mannheim, Germany). Cells transfected with mCitrine were imaged by excitation with the 514-nm laser line of a white light laser at 70% power with <10% transmission with 2.65 frames/s. Bleaching was performed by scanning a circular spot of 2 µm with 5 laser lines of the white light laser at 100% transmission (app. 1 mW per laserline) plus a 405-nm laser diode at 100% intensity. Cells were bleached 0.5, 2, 4, 6, 8, 10, 15 and 20 min after the beginning of fixation (compare fig. S1-S2). Fluorescence images were recorded before and after each bleaching. They were analysed using the software Fiji ^22^. To correct for total bleaching, development of autofluorescence and protein loss in the cell, the mean fluorescence intensity in the bleached spot was normalized to the mean fluorescence intensity of the whole cell. To correct for inhomogeneous distribution of fluorescence within the cell, it was further normalized to the initial ratio between the mean in bleaching spot (before the first bleaching) and the mean of the whole cell. The recovery of the normalized fluorescence between consecutive time points was calculated for every time point after the first one. As a control for fluorescence dark state recovery, cells that have been fixed for 1 h were bleached following the same protocol (Fig. S2). Fixation was therefore regarded as complete once the recovery of fluorescence intensity was no more significantly higher than its control. To evaluate completeness of protein fixation 1 h after onset of fixation, recovery curves over the first 20 s after a single bleaching were additionally recorded. To distinguish in this case between fluorescence dark state recovery and diffusion into the bleached spot, the bleached spot was divided into three concentric rings (Fig. S3). Diffusion into the spot is faster in the outer rings, whereas fluorescence dark state recovery is equally fast in all areas of the spot ^16^.

### Immunofluorescence microscopy

Immunofluorescence microscopy experiments were performed on HeLa or Cos7 cells grown in glass bottom dishes (MatTek Corporation, Ashland, MA, USA). Cells were fixed for 20 min at 37 °C either in 4% formaldehyde or 1% glutaraldehyde in 4% formaldehyde. After washing twice with PBS, cells were permeabelised with 0.5% Triton-X100 in PBS for 5 min. After washing twice with PBS again, the samples were incubated for 60 min with the primary antibodies in PBS containing 1% BSA. Alpha-tubulin was marked by a polyclonal antibody from rabbit (1:40; Abcam, Cambridge, UK). Antibodies for GM130 (1:200; BD Biosciences, Franklin Lakes, USA), HSP60 (1:100; Sigma-Aldrich Chemie GmbH, Munich, Germany) and LAMP2 (1:50; Fitzgerald industries Int., North Acton, MA, USA) were monoclonal mouse antibodies. PTP1b was marked with a polyclonal antibody from rabbit (1:200; Assay Biotechnology Company Inc., Fremont, CA, USA) and Rab5 with a monoclonal antibody from rabbit (Cell Signaling Technology Europe, Frankfurt, Germany). Labelling of Phalloidin-Alexa488 (1:200; Invitrogen Carlsbad, CA, USA) was done together with the PTP1b antibody. After washing twice with PBS again, the cells were incubated with the secondary antibodies (goat anti mouse coupled to Alexa 546 and goat anti rabbit coupled to Alexa 488; Thermo Fisher Scientific, Waltham, MA, USA) 1:300 in PBS containing 1% BSA. After washing twice with PBS again, DNA was stained by Hoechst 33342 (250 ng/mL in PBS) for 5 min. Before imaging cells were washed twice in PBS. Imaging was done on a Zeiss Axiovert widefield microscope using a 40x 1.3 NA or a 63x 1.4 NA oil immersion objective.

### Statistical analysis

Two-tailed, homoscedastic student’s t-tests were performed using the software Excel (Microsoft corporation, Redmond, WA, USA).

## Data availability

The data generated for this work are available from the corresponding author upon request.

## Acknowledgments

The authors would like to thank Michael Reichl for excellent technical assistance and Astrid Krämer for proofreading of the manuscript.

## Author Contributions

J.H. and B.K. designed and analyzed the experiments. J.H., J.S. and K.J.H. performed the experiments. J.H. wrote the manuscript.

## Competing financial interests

The authors declare that no competing interest exists.

